# Zidovudine multi-combos with last-line fosfomycin, ceftazidime-avibactam, colistin and tigecycline against Multi-Drug Resistant *Klebsiella pneumoniae*

**DOI:** 10.1101/2022.05.17.492182

**Authors:** Marta Gómara-Lomero, Ana Isabel López-Calleja, Antonio Rezusta, José Antonio Aínsa, Santiago Ramón-García

**Affiliations:** Department of Microbiology. Faculty of Medicine. University of Zaragoza, Spain; Servicio de Microbiología, Hospital Universitario Miguel Servet, Zaragoza, Spain; Research & Development Agency of Aragon (ARAID) Foundation, Spain; CIBER Respiratory Diseases, Carlos III Health Institute, Madrid, Spain

**Keywords:** antimicrobial resistance, MDR *Klebsiella pneumoniae*, synergy, fosfomycin, zidovudine, drug repurposing

## Abstract

Drug repurposing is a novel strategy for the development of new therapies against antibiotic-resistant bacteria. Zidovudine, an antiviral largely used in the HIV-therapy, exerts antibacterial activity against Gram-negative bacteria. Zidovudine was identified in a previous drug repurposing synergy screening as fosfomycin enhancer against *Klebsiella pneumoniae* ATCC 13883. Our aim was to evaluate the antibacterial *in vitro* activity of zidovudine-based combinations with last-line antibiotics against MDR/XDR *K. pneumoniae* isolates. We validated the zidovudine/fosfomycin combination against a collection of 12 MDR *K. pneumoniae* isolates by the checkerboard assay (CBA). In addition, we performed time-kill assays (TKA) to analyze synergistic and bactericidal activities of zidovudine paired combinations with fosfomycin, ceftazidime-avibactam, colistin and tigecycline. These were compared with frequent clinical combinations in the treatment of MDR Enterobacteriaceae. The potential of the triple zidovudine/fosfomycin/colistin was also assessed by TKA. CBA synergy confirmation rate between zidovudine/fosfomycin was 83.33%. TKA yielded synergy confirmation rates of 83.3% for zidovudine/ceftazidime-avibactam, 75% for zidovudine/fosfomycin, 75% for zidovudine/colistin and 66.6% for zidovudine/tigecycline with potent killing activities. Frequent clinical combinations displayed synergy rates of 41.6% for meropenem/ertapenem, 33.33% for meropenem/colistin, 75% for fosfomycin/colistin and 66.6% for fosfomycin/tigecycline with lower bactericidal efficacy than zidovudine-based combinations. The triple zidovudine/fosfomycin/colistin combination exhibited activities similar to fosfomycin/colistin and fosfomycin/zidovudine. As conclusion, zidovudine is an effective partner in *in vitro* combinations with existing antibiotics against MDR *K. pneumoniae*, especially with ceftazidime-avibactam, fosfomycin or colistin. Further studies are needed to elucidate the clinical potential of zidovudine as a repurposed drug in the antibacterial therapy.

## INTRODUCTION

The global prevalence and dissemination of MDR enterobacteria is facilitated by rapid acquisition of plasmid-mediated resistance mechanisms, including ESBL and carbapenemases that confer resistance to β-lactams, or the emergent *mcr-1* plasmid involved in colistin resistance (1). *Klebsiella pneumoniae* is the most concerning pathogen, leading the carbapenemase producing enterobacteria (CPE) and causing nosocomial infections with high mortality rates (2). The latest ECDC surveillance reported 7.9% carbapenem resistance and 19.3% combined resistance to traditional first-line antibiotics against *K. pneumoniae* in the EU (3), highlighting the complications in the therapeutic management of MDR-related infections.

Recommended treatments involve individualized combinatorial therapies including carbapenem-saving strategies and combinations of last-line agents such as colistin, tigecycline, aminoglycosides and fosfomycin (4,5). Few new antibiotics have been marketed lately. Novel molecules (e.g. cefiderocol, eravacycline) and β-lactam/β-lactamase inhibitor combinations (e.g. ceftazidime-avibactam, meropenem-vaborbactam) show good activities against MDR enterobacteria and are currently used in clinical practice (6–8). However, the emergence of resistant strains has been already reported (9–11), which compromise the use of these new agents that should be preserved for severe infections. New therapeutic strategies are thus urgently needed to improve efficacy of MDR-treatments and preserve the antibiotic arsenal.

Drug repurposing is an attractive strategy that allows faster clinical implementation, as pharmacokinetic and pharmacodynamic (PKPD) parameters and toxicity packages are already defined in commercialized drugs (12). The search for synergistic interactions has also emerged as a favorable approach to enhance the activity of drugs in combinatorial therapy. Moreover, this strategy has the potential to minimize toxicity and resistance emergence derived from monotherapy (13).

Zidovudine (3’-azido-3’-deoxythymidine), a thymidine analogue, was the first commercial antiretroviral agent for HIV/AIDS treatment. The antibacterial activity of zidovudine against Gram-negative bacteria is known since late 1980s (mainly due to the inhibition of bacterial DNA replication by targeting thymidine kinase (14)) demonstrating also *in vivo* efficacy (15,16). Toxicity and the emergence of resistant strains were the main drawbacks limiting the development of zidovudine-based antibacterial therapies. More recently, *in vitro* studies explored the synergistic activity of zidovudine in combination with known antibiotics (16–20).

In a previous work, we screened the FDA-library and identified zidovudine as a potent synergistic partner of fosfomycin against *K. pneumoniae* (*unpublished*). Here, we evaluated the synergistic and bactericidal *in vitro* activities of zidovudine in combination with fosfomycin, ceftazidime-avibactam, colistin and tigecycline against antibiotic-resistant *K. pneumoniae* isolates, and compared them with usual combinatorial treatments for MDR enterobacteria.

## MATERIALS AND METHODS

### Bacterial strains, clinical characterization and growth conditions

A well-characterized set of 12 MDR/XDR (21) *K. pneumoniae* isolates (eight from clinical samples and four from quality assessment exercises) was provided by the Miguel Servet University Hospital (Zaragoza, Spain), including representative resistance mechanisms (**Table S1**). Bacterial identification was performed by MALDI-TOF mass spectrometry (Bruker Daltonik GmbH, Germany) and antimicrobial susceptibility by an automated broth microdilution method (Microscan Walkaway®, Beckman Coulter, Spain). Phenotypic detection of ESBL, AmpC, carbapenemases and colistin resistance was done according to EUCAST guidelines (22). Genotypic characterization of resistance mechanisms was performed in clinical samples at the National Microbiology Centre (Majadahonda, Spain). Bacterial LB stocks (15% glycerol) were preserved at -20ºC. Freeze stocks were thawed and sub-cultured on Mueller Hinton broth for 24 hours at 37°C before each assay.

### Drugs susceptibility testing and media conditions

Zidovudine, fosfomycin disodium salt, glucose-6-phosphate, colistin sulfate, (Sigma–Aldrich, Darmstadt, Germany), tigecycline, ceftazidime (European Pharmacopoeia, Strasbourg, France), meropenem (Fresenius Kabi), ertapenem (MSD) and avibactam (AdooQ BioScience, Irvine, USA) were reconstituted in DMSO or water according to their solubilities. Stock solutions were prepared fresh on the same day of plate inoculation.

Drug susceptibility testing, checkerboard (CBA) and time-kill assays (TKA) were performed in CAMHB. MIC determinations were performed by broth microdilution in CAMHB following CLSI guidelines (23) linked to the MTT [3-(4,5-dimethylthiazol-2-yl)-2,5-diphenyl tetrazolium bromide] assay (24,25). Briefly, two-fold serial dilutions of drugs were inoculated with a bacterial suspension of 5×10^5^ CFU/mL in 96-well plates (V_F_= 150 μL) and incubated at 37°C for 18-20 hours. For fosfomycin and ceftazidime-avibactam susceptibility tests, CAMHB was supplemented with 25 mg/L of glucose-6-phosphate and 4 mg/L of avibactam, respectively (26). After incubation, 30 μL/well of a solution mix (MTT/Tween 80; 5 mg/mL/20%) were added and plates further incubated for 3 hours at 37ºC. MIC values were defined as the lowest concentration of drug that inhibited 90% of the OD_580_ MTT colour conversion (IC_90_) compared to growth control wells with no drug added.

MBC was also determined in order to discern bacteriostatic or bactericidal activities. Before MTT addition, 10 μL/well were transferred to 96-well plates containing LB agar and further incubated at 37ºC for 24 hours before addition of 30 μL/well of resazurin; a change from blue to pink indicated bacterial growth. The MBC was defined as the lowest concentration of drug that prevented this colour change. A compound was considered bactericidal when MBC/MIC ≤ 4 (24).

#### Synergy validation assays

##### (i) Checkerboard assays

Pairwise combinations of zidovudine and fosfomycin were assayed in 96-well plates using freshly prepared CAMHB. Each well was inoculated with 100 μl of a freshly grown bacterial suspension containing 5×10^5^ CFU/mL (V_F_=200 μL). Plates were incubated for 24 hours at 37°C and bacterial growth was measured using the MTT assay (24,25), as described above. Fractional Inhibitory Concentration Indexes (FICI) were calculated as the sum of FIC_ZDV_ plus FIC_FOF_; where FIC_ZDV_ is the MIC of zidovudine in the presence of fosfomycin divided by the MIC of zidovudine alone and, conversely, for FIC_FOF_. Synergy was defined as FICI ≤0.5, antagonism as FICI >4.0, and no interaction as FICI = 0.5 to 4.0 (27). The Fractional Bactericidal Concentration Index (FBCI) was similarly calculated based on MBC values, as described above.

##### (ii) Time-kill assays

Exponentially growing cultures of *K. pneumoniae* clinical strains were inoculated in duplicates in CAMHB 96-well plates (V_F_= 280 μL/well; 5×10^5^ CFU/mL) containing increasing concentrations (0.1x, 0.25x, 1x, 4x, 10xMIC values) of compounds alone, and incubated at 37ºC. Drug-free wells were used as growth controls and MIC assays were performed in parallel with the same inoculum to ensure compound activity. Samples were taken at 0, 2, 5, 8, 24 and 48 hours, and bacterial population was quantified by spot-platting 10-fold serial dilutions onto MHA plates. Plates were incubated overnight at 37°C and CFU/mL calculated. The lower limit of detection was 50 CFU/mL. To assess the activity of the combinations, generated dose-response curves of compounds alone were analyzed to select appropriate test combo concentrations (up to 300 mg/L if compound was inactive) and TKA equally performed as described above. Zidovudine combinations were also tested at concentrations of 1 mg/L for the twelve clinical strains, including those showing high-level zidovudine resistance, to assess physiological relevant concentrations, i.e. C_max_ of zidovudine observed in human plasma after intravenously administration (1.1 to 1.8 mg/L) (28).

The activity of the four novel zidovudine-based combinations described in this work (fosfomycin/zidovudine, ceftazidime-avibactam/zidovudine, colistin/zidovudine and tigecycline/zidovudine) was compared with that of four usual MDR-treatments (meropenem/ertapenem, meropenem/colistin, fosfomycin/colistin and fosfomycin/tigecycline). The triple fosfomycin/colistin/zidovudine combination was also tested against eight strains and compared with the activity of the three drugs alone and in pairwise combinations at matching concentration (0.25-1xMIC). Synergistic and bactericidal activities were evaluated after 8, 24 and 48 hours of incubation. In TKA, synergy was defined when there was a ≥2 log_10_ CFU/mL decrease in bacterial count compared to the most active single agent in the combination at any time point (8, 24 and 48 hours). Antagonism was defined as a ≥2 log_10_ increase in CFU/mL between the combination and the most active single agent. All other degrees of interaction were defined as indifferent. Bactericidal activity was defined as a ≥3 log_10_ CFU/mL reduction at 8, 24 and 48 hours compared to the initial inoculum (29).

## RESULTS

We followed a stepwise approach in order to characterize the potential of zidovudine in the treatment of infections caused by MDR *K. pneumoniae*: first, the activity of test compound was determined by drug susceptibility assays; second, pairwise and triple combinations were tested by CBA and TKA to identify those most promising synergistic combinations. All these experiments were performed against a panel of twelve MDR/XDR *K. pneumoniae* isolates with representative mechanisms of resistance.

### Drug susceptibility characterization

Zidovudine MIC values ranged from 0.25 to ≥64 mg/L, with nine strains showing low MIC_ZDV_ values (0.25 to 2 mg/L) and three other strains showing MIC values ≥16 mg/L; in consequence, 2 mg/L could be a possible cut-off for zidovudine resistance. The number of multidrug-resistant determinants appeared not to be related with the MIC_ZDV_ values. The susceptibility profiles obtained for the drugs tested among the twelve MDR/XDR *K. pneumoniae* isolates according to EUCAST breakpoints (26) are also shown **Table S1**.

### Zidovudine/fosfomycin combinations displayed wide coverage synergy against MDR/XDR *K. pneumoniae* isolates

Synergy between fosfomycin and zidovudine was confirmed in 75% (9/12) and 66.6% (8/12) of the strains by FICI and FBCI, respectively. Interestingly, effective fosfomycin concentrations in the presence of zidovudine were reduced 2-16-fold, restoring fosfomycin susceptibility below the EUCAST breakpoint (MIC≤32 mg/L) in all strains. Antimicrobial zidovudine concentrations were also significantly lower in presence of fosfomycin, with reductions ranging from 2- to 128-fold, and ≥4-fold reduction in 75% (9/12) and 90.9% (10/11) of the strains by FICI and FBCI, respectively. No antagonisms were observed. (**Table 1**).

**Table 1.** Pairwise interactions of fosfomycin plus zidovudine against *K. pneumoniae* isolates. The CBA was used to evaluate the degree of interaction. FICI, Fractional Inhibitory Concentration Index; FBCI, Fractional Bactericidal Concentration Index. Values in bold indicate synergy (FICI, FBCI ≤0.5), while values >0.5 indicate no interaction. FOF, fosfomycin; ZDV, zidovudine.

| Isolate | MIC (mg/L) alone |  | MIC (mg/L) in combination |  | MBC (mg/L) alone |  | MBC (mg/L) in combination |  | FICI | FBCI |
| --- | --- | --- | --- | --- | --- | --- | --- | --- | --- | --- |
|  | FOF | ZDV | FOF | ZDV | FOF | ZDV | FOF | ZDV |  |  |
| E-1 | >64 | 0.5 | 16 | 0.0625 | >64 | 2 | 32 | 0.5 | **0.38** | 0.75 |
| E-2 | >64 | >4 | 8 | 1 | >64 | >4 | 8 | 1 | **0.19** | **0.19** |
| E-3 | >64 | 2 | 32 | 1 | >64 | 2 | 32 | 1 | 0.75 | 0.75 |
| E-4 | 64 | 1 | 16 | 0.25 | 64 | >2 | 16 | 0.5 | **0.50** | **0.38** |
| E-5 | >16 | 2 | 4 | 0.25 | >16 | >2 | 4 | 0.25 | **0.25** | **0.19** |
| A-6 | >64 | 16 | 8 | 0.5 | >64 | 16 | 8 | 1 | **0.09** | **0.13** |
| C-7 | >64 | 1 | 16 | 0.125 | >64 | 2 | 16 | 0.125 | **0.25** | **0.19** |
| CS-8 | 64 | 4 | 16 | 1 | 64 | 4 | 32 | 2 | **0.5** | 0.75 |
| CSE-9 | >64 | $\geq 64$ | 16 | 1 | >64 | $\geq 64$ | 16 | 1 | **0.13** | **0.13** |
| CE-10 | >64 | 64 | 8 | 0.5 | >64 | >64 | 8 | 2 | **0.07** | **0.08** |
| CEE-11 | >64 | 1 | 16 | 0.5 | >64 | >4 | 16 | 2 | 0.63 | **0.38** |
| CSEE-12 | 64 | 8 | 16 | 4 | 128 | 8 | 64 | 0.25 | 1 | 0.53 |

### Zidovudine-based combinations are more potent *in vitro* than current clinically used combinations against MDR/XDR *K. pneumoniae* isolates

The use of CBA allows screening of a number of antibiotic combinations in search of synergy, although it is limited to single-time point read-outs. In order to give robustness to the interaction analysis of zidovudine-based combinations, we performed TKA that provide more detailed synergistic information, including bactericidal and sterilizing activities of the combinations over time (**Figure 1**). At any time point (8, 24 and 48 hours), synergy rates among currently used combinations for MDR treatment was observed in several isolates: 41.6% for meropenem/ertapenem (5/12), 33.33% for meropenem/colistin (4/12), 75% for fosfomycin/colistin (9/12) and 66.6% for fosfomycin/tigecycline (8/12) (**Figure S1**). Interestingly, among all isolates, the highest number of synergistic interactions were obtained with zidovudine-based combinations, at zidovudine concentrations ranging from 0.004x to 2xMIC (**Figure S2**). The combination of ceftazidime-avibactam plus zidovudine was the most active, showing a synergy rate among the isolates of 83.3% (10/12). In 8 out of 12 strains the killing activity was below the limit of detection after 24 hours of incubation, preventing bacterial regrowth (a proxy for sterilizing activity) (**Figure S2a**). The combination of fosfomycin plus zidovudine was synergistic in 9 out of 12 strains, showing a potent and rapid initial decrease in bacterial counts after 8 hours in five strains (E-5, A-6, C-7, CSE-9, CE-10) and bactericidal activity down to the limit of detection in six strains (E-5, A-6, C-7, CSE-9, CEE-11 and CS-8) (**Figure S2b**). The combination of zidovudine plus colistin had 75% (9/12) synergy rates and 33.3% (4/12) killing rates down to the limit of detection (**Figure S2c**). Finally, the combination of zidovudine plus tigecycline was the less potent one showing late synergy (48 hours) in 66.6% (8/12) of the strains with bactericidal activity only against C-7 (**Figure S2d**).

**Figure 1.**
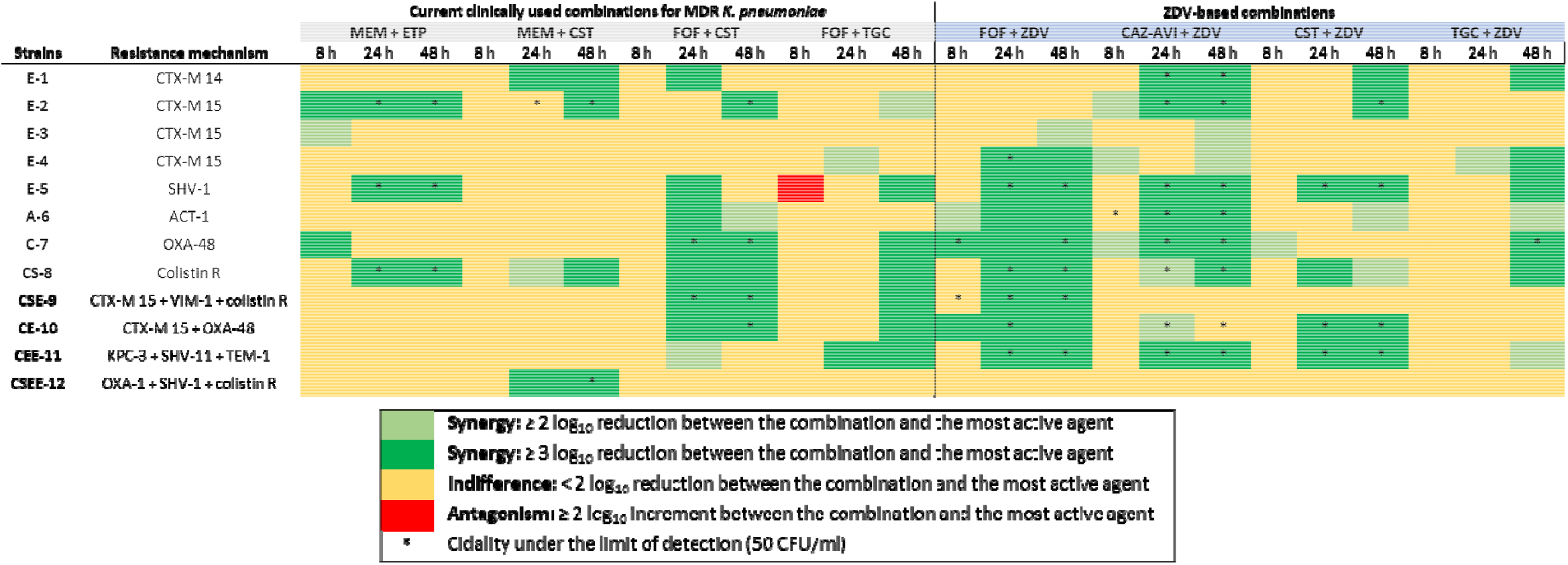
Heat map representation of synergy and bactericidal activities in pairwise combinations at different time points obtained by time-kill assays against *K. pneumoniae* isolates. When several concentrations were tested for the same drug, the most favourable outcome is displayed. ZDV-based combinations were tested at concentrations ≤1 mg/L, reflecting physiologically achieved concentrations. Data supporting this summary figure is displayed in Figure S1 and Figure S2. CAZ-AVI; ceftazidime-avibactam; CST, colistin; ETP, ertapenem; FOF, fosfomycin; MEM, meropenem; TGC, tigecycline; ZDV, zidovudine.

Notably, the synergistic killing effects of zidovudine-based combinations were observed even at low zidovudine concentrations (≤1 mg/L), which are below pharmacological serum concentrations at standard zidovudine oral doses. This effect was observed regardless the zidovudine susceptibility profile of the strains (A-6, CSE-9 and CE-10 are zidovudine-resistant) (**Figure 3)**.

### Triple zidovudine-based combination offers limited advantages over already synergistic pairwise combinations

We identified that while carbapenem-based combinations had little synergistic interaction profiles, the combination of two last-line antibiotics (fosfomycin/colistin) had an overall synergy rate of 75%. Interestingly, zidovudine displayed strong synergy with both drugs. We thus aimed to characterize whether the addition of zidovudine to the fosfomycin/colistin combination could further potentiate the synergistic interaction, as previously described (24,30,31). The triple combination was highly active showing bactericidal activity to the limit of detection in 4 out of 8 strains tested. However, compared to the fosfomycin/zidovudine, colistin/zidovudine and fosfomycin/colistin combinations, fosfomycin/colistin/zidovudine was more effective at bacterial eradication than colistin/zidovudine, but added little efficacy when compared to fosfomycin/zidovudine or fosfomycin/colistin (**Figures 2 & 4)**.

**Figure 2.**
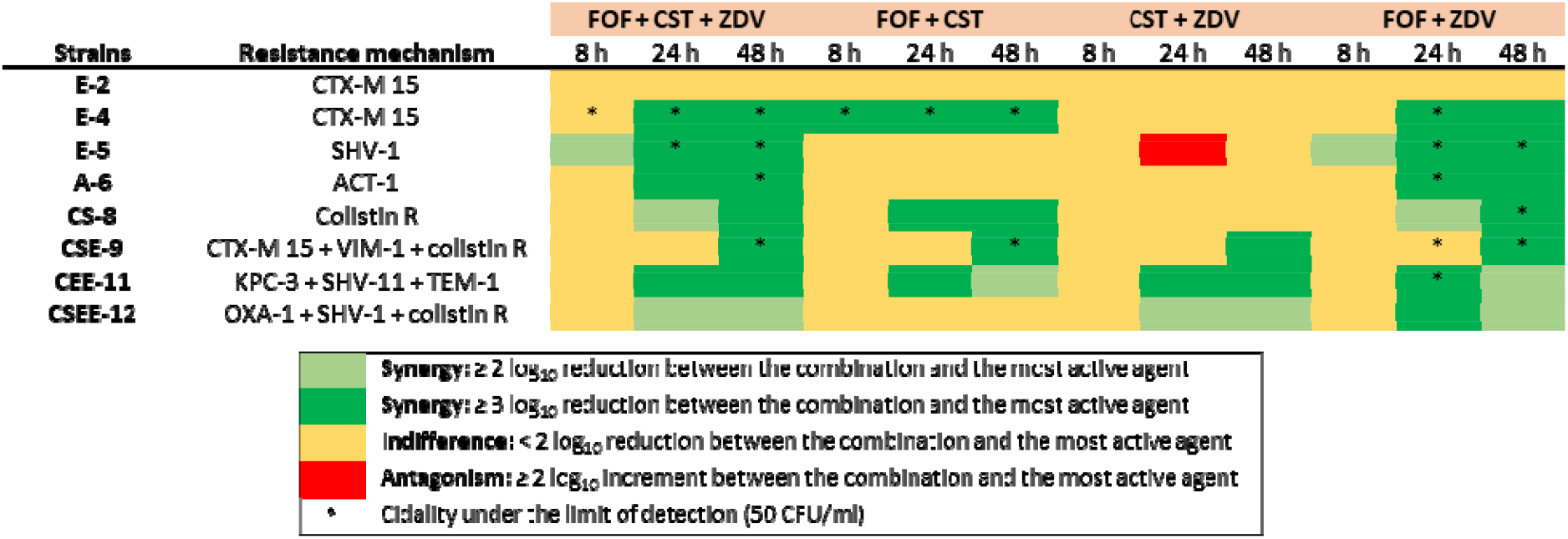
Heat map representation of synergy and bactericidal activities in triple combination (fosfomycin/colistin/zidovudine) compared to those in pairwise combinations at different time points obtained by time-kill assays against eight *K. pneumoniae* isolates. Combo tested concentrations at 0.25-1x MIC. Data supporting this summary figure is displayed in Figure 4. CST, colistin; FOF, fosfomycin; ZDV, zidovudine.

**Figure 3.**
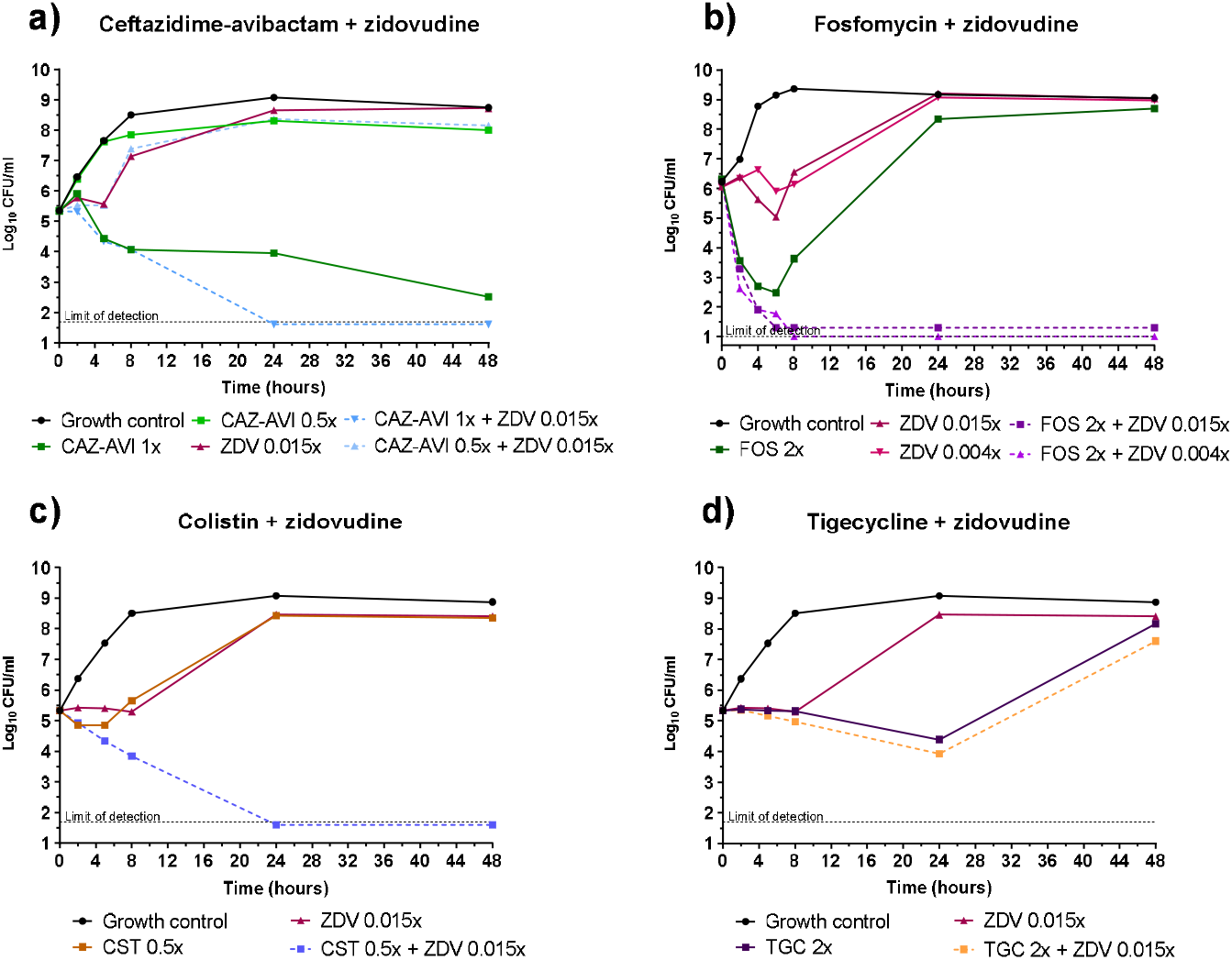
Time–kill assays characterization of zidovudine combinations with last-line antibiotics in the treatment of MDR-*K. pneumoniae* infections. The zidovudine resistant clinical strain CE-10 (*bla*_OXA-48_ + *bla*_CTX-M15_) was used for these studies. Zidovudine enhanced the activities of ceftazidime-avibactam, fosfomycin and colistin even at low sub-inhibitory concentrations (0.004-0.015x MIC), showing potent synergistic and bactericidal effects with last-line antibiotics. ZDV concentrations were 1 mg/L, which is below C_max_ values (1.1 to 1.8 mg/L) achieve in human plasma after a recommended ZDV oral dose. MIC_CAZ-AVI_= 0,25 mg/L, MIC_CST_= 1 mg/L, MIC_FOF_= 64 mg/L, MIC_TGC_= 1 mg/L, MIC_ZDV_= 64 mg/L.

**Figure 4.**
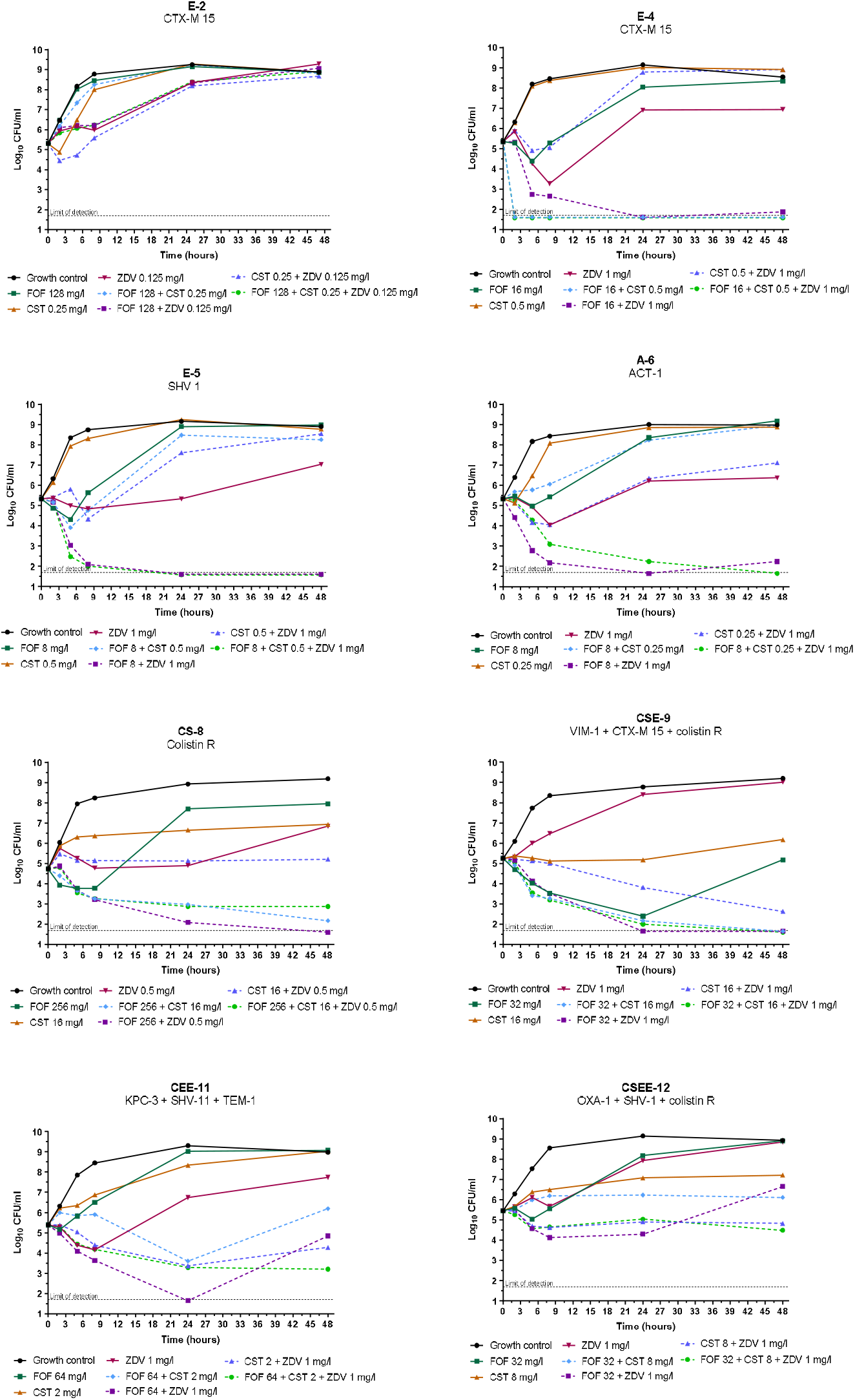
Time-kill curves of zidovudine, colistin and fosfomycin alone, pairwise and triple combination against eight K. pneumoniae isolates.

## DISCUSSION

We evaluated the *in vitro* efficacy of zidovudine in combination with antibiotics currently used for the treatment of infections caused by MDR/XDR *K. pneumoniae* with common resistance patterns, such as ceftazidime-avibactam, colistin, fosfomycin and tigecycline. For this, we used a panel of 12 MDR/XDR *K. pneumoniae* isolates and tested zidovudine at clinically achievable concentrations. Our TKA data showed high rates of synergistic and killing activities of zidovudine-based combinations even against strains with concurrent resistance mechanisms to these and other antimicrobials, suggesting a potential role of zidovudine in combinatorial therapy (**Figures 1, S1 & S2**).

We performed extensive TKA and obtained readouts after up to 48 hours of incubation. We first characterized the activity of the compounds alone in a dose-response manner against all isolates. Then, we selected subinhibitory/static concentrations that were matched for the combination assays to allow for a wider dynamic range; using higher/effective concentrations of the compounds alone would mask the effect of any potential interaction. Our efficacy analysis then takes into consideration, not only the increased bactericidal activity of the combination compared to the drugs alone, but also the ability of the combination to completely eradicate bacteria down to the limit of detection of the assay (a proxy for sterilization of the culture). Based on these criteria, we tested several zidovudine-based combinations and compared them with current combinations clinically used to treat MDR/XDR *K. pneumoniae* infections (**Figure 1**).

### (i) Zidovudine plus ceftazidime-avibactam

This combination showed the highest efficacy with potent killing activity in all except two strains (CSE-9 and CSEE-12) even at low concentrations of ceftazidime-avibactam (ranging from 0.125x to 1xMIC) (**Figures 1 & S2a**). CSE-9 displayed high MIC values to ceftazidime-avibactam (>64 mg/l) and zidovudine (≥64 mg/L), while for CSEE-12 these were low (MIC_CAZ-AVI_= 0.5 mg/L, MIC_ZDV_= 1 mg/L); further studies are needed to elucidate whether acquired resistance mechanisms might explain this lack of activity.

Safety and efficacy of ceftazidime-avibactam against MDR enterobacteria facilitated its inclusion as first-line therapeutic option for infections caused by CPE. It is administered in monotherapy against OXA-48 (class D) and KPC (class A) producers or associated to aztreonam against class B enzymes (β-lactamases refractory to inhibition by avibactam) (32). Although the potential for resistance selection appears to be low (33), the extensive use of ceftazidime-avibactam as a savage therapy will contribute to the emergence of resistance. In fact, resistance linked to mutations in plasmid-borne KPC-3 were reported during ceftazidime-avibactam treatment (34,35), and development of resistance to ceftazidime-avibactam is more likely after previous exposure with meropenem-vaborbactam (36,37).

To the best of our knowledge, this is the first time that the zidovudine/ceftazidime-avibactam combination is evaluated against MDR *K. pneumoniae*. Our promising results might lay the foundations of further studies to support a potential clinical implementation.

### (ii) Zidovudine plus fosfomycin

We identified zidovudine/fosfomycin synergy in a previous screening (*unpublished*). Our data were in agreement with Antonello *et al*. describing such synergy by CBA in 69.4% of Enterobacteriaceae strains, including fosfomycin-resistant strains, and characterized its killing effects by TKA at 24 hours (17). In our work, we further validate this synergism (**Figure 1**); fosfomycin alone showed a fast bactericidal activity followed by a sharp growth rebound and, although strains in our collection displayed high MIC_FOF_ values (from 8 to ≥64 mg/L), the combination with zidovudine was able to restore fosfomycin activity, preventing this bacterial regrowth (**Figure S2b**).

### (iii) Zidovudine plus colistin

Our interaction data is also in agreement with previously reported (16,19), including *K. pneumoniae* colistin-resistant strains (20). However, TKA against our colistin-resistant strains revealed both synergy against CS-8 (MIC_CST_= 16 mg/L) (but a lack of bactericidal activity) and lack of interaction against CSE-9 and CSEE-12 (MIC_CST_= 16 and 4 mg/L, respectively) with concurrent alternative resistance mechanisms (**Figure S2c**).

### (iv) Zidovudine plus tigecycline

This was the least effective combination (similar to fosfomycin/tigecycline) with just late synergy and bacteriostatic profile at 48 hours. The activities were also tigecycline dose-dependent for most strains (**Figures S1d and S2d**). Nevertheless, TKA were able to revealed important interactions confronting data generated by CBA (18). The fact that zidovudine/tigecycline showed a lower degree of interaction could be explained because both drugs have to reach their intracellular targets; however, in other combos, when zidovudine is used along with extracellular targeting compounds such as fosfomycin, ceftazidime-avibactam and colistin, the action of the latter drugs could result in an increased permeability of zidovudine, hence resulting in a higher effectivity (16,19,20,38). Adding to this hypothesis, the use of glucose-6-phosphate (added in *in vitro* experiments to mimic physiological conditions and promote intracellular transport of fosfomycin) might also facilitate zidovudine entrance to the bacterial cell (17).

### (v) Zidovudine plus fosfomycin/colistin

The synergism between two last-line antibiotics fosfomycin/colistin against MDR *K. pneumoniae* was previously reported by TKA and hollow-fiber infection model (39,40), and clinical studies (41,42). We also identified such a positive interaction between fosfomycin/colistin, two drugs that displayed independently potent activities in combination with zidovudine (**Figure 1**). Given that our studies demonstrated that zidovudine-based combinations are more potent than currently clinically used for the treatment of infections caused by MDR/XDR *K. pneumoniae* strains, we performed TKA to characterize the potential impact of zidovudine in a triple combination with fosfomycin and colistin. In mycobacteria, triple combinations perform better than the sum of the pairwise combinations (24,30,31); however, this was not the case and the triple combination added little value to fosfomycin/colistin (**Figure 2**).

For HIV treatment, zidovudine is dosed at 500-600 mg daily oral and 1.5 mg/kg/6h intravenously. After standard dosage regimens, C_max_ ranging from 1.1 to 1.8 mg/L are achieved (28,43). Previous i*n vitro* and *in vivo* studies suggested that clinically achievable zidovudine concentrations could be effective against MDR enterobacteria when in combination therapy (16,17,20). Zidovudine toxicity is associated to the dose, disease stage and prolonged HIV-therapy. Safety profiles observed in HIV-patients together with a short plasma half-life (1.1–2.3 hours) (28,43,44) suggest that appearance of zidovudine toxicities are unlikely. Most reported side effects include headaches, myalgia, nausea and vomiting. Major toxicities (anemia and neutropenia) are more frequently described at high doses (1.200-1.500 mg/day) after more than 4 weeks of treatment (45,46). A few case reports described oral zidovudine overdoses up to 36 g/daily without abnormalities or with slight and transient side-effects such as lethargy (47).

In our study, MICs of zidovudine ranged from 0.25 to ≥64 mg/L, which are in accordance with similar studies (16,17,19,20). We observed potent bactericidal activities of the combinations against most strains at zidovudine concentrations below 1 mg/L with the exception of A-6 strain for which the effective zidovudine concentration in combinations was 4 mg/ml (**Figure S2**), which still would be below C_max_ expected at daily doses of 600 mg (17). Pharmacokinetics and safety of zidovudine plus colistin combination antimicrobial therapy was evaluated in a clinical trial (48,49). It was found that doses of both synergistic partners, zidovudine and colistin (which has toxicity issues), could be reduced while retaining their therapeutic efficacy (16). Our data thus suggests that current zidovudine dosing strategies might suffice to treat bacterial infections in humans and that zidovudine associated side effects are unlikely to occur during short-term regimens, as in the context of acute bacterial infections. In addition, zidovudine reduces transmission of ESBL and carbapenemase containing plasmids, hence supporting zidovudine use in the prevention of the spread of resistant enterobacteria (50).

Future directions will include: (i) expanding such studies to a larger panel of clinical isolates from several locations; (ii) identifying additional resistance mechanisms (i.e. porin loss or efflux pumps) besides the genotypic characterization of our strain panel that included standard β-lactam enzymatic resistance; and (iii) investigating for deficiency in thymidine kinase genes (which normally phosphorylate inactive zidovudine into the active form (51,52)) in those strains exhibiting high MIC_ZDV_ values, since it remains unknown whether the selection of resistant mutants is responsible for the bacterial rebound observed by TKA in some combinations (**Figure S2**).

In conclusion, zidovudine in combination with other antimicrobial drugs is a repurposing option for MDR/XDR *K. pneumoniae*; similar repurposing approaches that employ other nucleoside analogues in combination with antifungals are already in clinical use (53). Based on our studies, we propose the following priority list of pairwise combinations: zidovudine/ceftazidime-avibactam > zidovudine/fosfomycin > fosfomycin/colistin > zidovudine/colistin > fosfomycin/tigecycline = zidovudine/tigecycline > meropenem/colistin > meropenem/ertapenem. Finally, further dynamic PKPD studies would be needed to fully discern the potential of zidovudine-based combinations in the clinical practice, and future clinical trials would clarify the impact of zidovudine combination therapy on clinical outcomes. If these studies result in clinical improvement, future expectations could include individualised therapy in those patients with severe infections. As part of routine laboratory workflow, synergy testing with zidovudine in clinical isolates would allow to infer the success of combination therapy.

## Data availability statement

All data pertaining to this work is within the main manuscript or supplementary information.

## REFERENCES

1. Chew KL, Lin RTP, Teo JWP. Klebsiella pneumoniae in Singapore: Hypervirulent infections and the carbapenemase threat. Front Cell Infect Microbiol. 2017;7:515.

2. Lee CR, Lee JH, Park KS, Kim YB, Jeong BC, Lee SH. Global dissemination of carbapenemase-producing Klebsiella pneumoniae: Epidemiology, genetic context, treatment options, and detection methods. Front Microbiol. 2016;7:895.

3. European Centre for Disease Prevention and Control. Antimicrobial resistance in the EU/EEA (EARS-Net), Annual Epidemiological Report for 2019. ECDC. Stockholm; 2020.

4. Bassetti M, Peghin M, Pecori D. The management of multidrug-resistant Enterobacteriaceae. Curr Opin Infect Dis. 2016 Dec;29(6):583–94.

5. Karaiskos I, Antoniadou A, Giamarellou H. Combination therapy for extensively-drug resistant gram-negative bacteria. Expert Rev Anti Infect Ther. 2017;15(12):1123–40.

6. Pogue JM, Bonomo RA, Kaye KS. Ceftazidime/Avibactam, Meropenem/Vaborbactam, or Both? Clinical and Formulary Considerations. Clin Infect Dis. 2019 Jan 18;68(3):519–24.

7. Wu JY, Srinivas P, Pogue JM. Cefiderocol: A Novel Agent for the Management of Multidrug-Resistant Gram-Negative Organisms. Infect Dis Ther. 2020 Mar;9(1):17–40.

8. Johnston BD, Thuras P, Porter SB, Anacker M, VonBank B, Vagnone PS, et al. Activity of cefiderocol, ceftazidime-avibactam, and eravacycline against carbapenem-resistant Escherichia coli isolates from the United States and International sites in relation to clonal background, resistance genes, coresistance, and region. Antimicrob Agents Chemother. 2020 Sep 21;64(10):e00797–20.

9. Zhong H, Zhao XY, Zhang ZL, Gu ZC, Zhang C, Gao Y, et al. Evaluation of the efficacy and safety of ceftazidime/avibactam in the treatment of Gram-negative bacterial infections: a systematic review and meta-analysis. Int J Antimicrob Agents. 2018;52(4):443–50.

10. Wang L, Liu D, Lv Y, Cui L, Li Y, Li T, et al. Novel Plasmid-Mediated tet(X5) Gene Conferring Resistance to Tigecycline, Eravacycline, and Omadacycline in a Clinical Acinetobacter baumannii Isolate. Antimicrob Agents Chemother. 2020 Dec 20;64(1):e01326–19.

11. Dulyayangkul P, Ismah Wakwn, Douglas EJA, Avison MB. Mutation of kvrA causes OmpK35 and OmpK36 porin downregulation and reduced meropenem-vaborbactam susceptibility in KPC-Producing Klebsiella pneumoniae. Antimicrob Agents Chemother. 2020 Jun 23;64(7):e02208–19.

12. Pushpakom S, Iorio F, Eyers PA, Escott KJ, Hopper S, Wells A, et al. Drug repurposing: progress, challenges and recommendations. Nat Rev Drug Discov. 2019;18(1):41–58.

13. Sun W, Sanderson PE, Zheng W. Drug combination therapy increases successful drug repositioning. Drug Discov Today. 2016 Jul;21(7):1189–95.

14. Lewin CS, Amyes SG. Conditions required for the antibacterial activity of zidovudine. Eur J Clin Microbiol Infect Dis. 1989;8(8):737–41.

15. Keith BR, White G, Wilson HR. In vivo efficacy of zidovudine (3’-azido-3’-deoxythymidine) in experimental gram-negative-bacterial infections. Antimicrob Agents Chemother. 1989;33(4):479–83.

16. Hu Y, Liu Y, Coates A. Azidothymidine produces synergistic activity in combination with colistin against antibiotic-resistant Enterobacteriaceae. Antimicrob Agents Chemother. 2018;63(1):e01630–18.

17. Antonello RM, Di Bella S, Betts J, La Ragione R, Bressan R, Principe L, et al. Zidovudine in synergistic combination with fosfomycin: an in vitro and in vivo evaluation against multidrug-resistant Enterobacterales. Int J Antimicrob Agents. 2021;58(1):106362.

18. Ng SMS, Sioson JSP, Yap JM, Ng FM, Ching HSV, Teo JWP, et al. Repurposing Zidovudine in combination with Tigecycline for treating carbapenem-resistant Enterobacteriaceae infections. Eur J Clin Microbiol Infect Dis. 2018 Jan;37(1):141–8.

19. Lin YW, Rahim NA, Zhao J, Han ML, Yu HH, Wickremasinghe H, et al. Novel polymyxin combination with the antiretroviral zidovudine exerts synergistic killing against NDM-producing multidrug-resistant Klebsiella pneumoniae. Antimicrob Agents Chemother. 2019 Mar 27;63(4):e02176–18.

20. Falagas ME, Voulgaris GL, Tryfinopoulou K, Giakkoupi P, Kyriakidou M, Vatopoulos A, et al. Synergistic activity of colistin with azidothymidine against colistin-resistant Klebsiella pneumoniae clinical isolates collected from inpatients in Greek hospitals. Int J Antimicrob Agents. 2019 Jun;53(6):855–8.

21. Magiorakos A-P, Srinivasan A, Carey RB, Carmeli Y, Falagas ME, Giske CG, et al. Multidrug-resistant, extensively drug-resistant and pandrug-resistant bacteria: an international expert proposal for interim standard definitions for acquired resistance. Clin Microbiol Infect. 2012 Mar;18(3):268–81.

22. EUCAST. The European Committee on Antimicrobial Susceptibility Testing. EUCAST guidelines for detection of resistance mechanisms and specific resistances of clinical and / or epidemiological importance. Version 2.0. 2017.

23. CLSI. Performance Standards for Antimicrobial Susceptibility Testing: 27th edition. CLSI supplement M100. Wayne, PA: Clinical and Laboratory Standards Institute; 2017.

24. Ramón-García S, Ng C, Anderson H, Chao JD, Zheng X, Pfeifer T, et al. Synergistic drug combinations for tuberculosis therapy identified by a novel high-throughput screen. Antimicrob Agents Chemother. 2011;55(8):3861–9.

25. Montoro E, Lemus D, Echemendia M, Martin A, Portaels F, Palomino JC. Comparative evaluation of the nitrate reduction assay, the MTT test, and the resazurin microtitre assay for drug susceptibility testing of clinical isolates of Mycobacterium tuberculosis. J Antimicrob Chemother. 2005 Apr;55(4):500–5.

26. EUCAST. The European Committee on Antimicrobial Susceptibility Testing. Breakpoint tables for interpretation of MICs and zone diameters. Version 11.0. 2021.

27. Odds FC. Synergy, antagonism, and what the chequerboard puts between them. J Antimicrob Chemother. 2003;52(1):1.

28. Wei L, Mansoor N, Khan RA, Czejka M, Ahmad T, Ahmed M, et al. WB-PBPK approach in predicting zidovudine pharmacokinetics in preterm neonates. Biopharm Drug Dispos. 2019;40(9):341–9.

29. Eliopoulos GM, Moellering RC. Antimicrobial combinations. In: Lorian V, editor. Antibiotics in laboratory medicine. 4th ed. Baltimore, MD: The Williams & Wilkins Co.,; 1996. p. 330–96.

30. Arenaz-Callao MP, González del Río R, Lucía Quintana A, Thompson CJ, Mendoza-Losana A, Ramón-García S. Triple oral beta-lactam containing therapy for Buruli ulcer treatment shortening. PLoS Negl Trop Dis. 2019;13(1):e0007126.

31. Ramón-García S, González Del Río R, Villarejo AS, Sweet GD, Cunningham F, Barros D, et al. Repurposing clinically approved cephalosporins for tuberculosis therapy. Sci Rep. 2016;6:34293.

32. Tamma PD, Aitken SL, Bonomo RA, Mathers AJ, van Duin D, Clancy CJ. Infectious Diseases Society of America Guidance on the Treatment of Extended-Spectrum β-lactamase Producing Enterobacterales (ESBL-E), Carbapenem-Resistant Enterobacterales (CRE), and Pseudomonas aeruginosa with Difficult-to-Treat Resistance (DTR-P. aerug. Clin Infect Dis. 2021;72(7):e169–83.

33. Shirley M. Ceftazidime-Avibactam: A Review in the Treatment of Serious Gram-Negative Bacterial Infections. Drugs. 2018 Apr;78(6):675–92.

34. Haidar G, Clancy CJ, Shields RK, Hao B, Cheng S, Nguyen MH. Mutations in blaKPC-3 That Confer Ceftazidime-Avibactam Resistance Encode Novel KPC-3 Variants That Function as Extended-Spectrum β-Lactamases. Antimicrob Agents Chemother. 2017 Apr;61(5):e02534–16.

35. Shields RK, Chen L, Cheng S, Chavda KD, Press EG, Snyder A, et al. Emergence of Ceftazidime-Avibactam Resistance Due to Plasmid-Borne blaKPC-3 Mutations during Treatment of Carbapenem-Resistant Klebsiella pneumoniae Infections. Antimicrob Agents Chemother. 2017 Feb;61(3):e02097–16.

36. Ackley R, Roshdy D, Meredith J, Minor S, Anderson WE, Capraro GA, et al. Meropenem-Vaborbactam versus Ceftazidime-Avibactam for Treatment of Carbapenem-Resistant Enterobacteriaceae Infections. Antimicrob Agents Chemother. 2020 Apr 21;64(5):e02313–19.

37. Shields RK, Chen L, Cheng S, Chavda KD, Press EG, Snyder A, et al. Emergence of Ceftazidime-Avibactam Resistance Due to Plasmid-Borne bla KPC-3 Mutations during Treatment of Carbapenem-Resistant Klebsiella pneumoniae Infections. Antimicrob Agents Chemother. 2017 Mar;61(3):e02097–16.

38. Antonello RM, Principe L, Maraolo AE, Viaggi V, Pol R, Fabbiani M, et al. Fosfomycin as partner drug for systemic infection management. A systematic review of its synergistic properties from in vitro and in vivo studies. Antibiotics. 2020;9(8):500.

39. Zhao M, Bulman ZP, Lenhard JR, Satlin MJ, Kreiswirth BN, Walsh TJ, et al. Pharmacodynamics of colistin and fosfomycin: A “treasure trove” combination combats KPC-producing Klebsiella pneumoniae. J Antimicrob Chemother. 2017;72(7):1985–90.

40. Wang J, He JT, Bai Y, Wang R, Cai Y. Synergistic activity of colistin/fosfomycin combination against carbapenemase-producing Klebsiella pneumoniae in an in vitro pharmacokinetic/pharmacodynamic model. Biomed Res Int. 2018 Apr 23;2018:5720417.

41. Michalopoulos A, Virtzili S, Rafailidis P, Chalevelakis G, Damala M, Falagas ME. Intravenous fosfomycin for the treatment of nosocomial infections caused by carbapenem-resistant Klebsiella pneumoniae in critically ill patients: a prospective evaluation. Clin Microbiol Infect. 2010 Feb;16(2):184–6.

42. Pontikis K, Karaiskos I, Bastani S, Dimopoulos G, Kalogirou M, Katsiari M, et al. Outcomes of critically ill intensive care unit patients treated with fosfomycin for infections due to pandrug-resistant and extensively drug-resistant carbapenemase-producing Gram-negative bacteria. Int J Antimicrob Agents. 2014 Jan;43(1):52–9.

43. Fillekes Q, Kendall L, Kitaka S, Mugyenyi P, Musoke P, Ndigendawani M, et al. Pharmacokinetics of zidovudine dosed twice daily according to World Health Organization weight bands in Ugandan HIV-infected children. Pediatr Infect Dis J. 2014;33(5):495–8.

44. Moore KHP, Raasch RH, Brouwer KLR, Opheim K, Cheeseman SH, Eyster E, et al. Pharmacokinetics and bioavailability of zidovudine and its glucuronidated metabolite in patients with human immunodeficiency virus infection and hepatic disease (AIDS clinical trials group protocol 062). Antimicrob Agents Chemother. 1995;39(12):2732–7.

45. Rachlis A, Fanning MM. Zidovudine Toxicity: Clinical Features and Management. Drug Saf. 1993;8(4):312–20.

46. McLeod GX, Hammer SM. Zidovudine: Five Years Later. Ann Intern Med. 1992 Sep 15;117(6):487–501.

47. Kroon S, Worm AM. Zidovudine overdose. Int J STD AIDS. 1991;2(1):56–7.

48. Loose M, Naber KG, Hu Y, Coates A, Wagenlehner FME. Serum bactericidal activity of colistin and azidothymidine combinations against mcr-1-positive colistin-resistant Escherichia coli. Int J Antimicrob Agents. 2018 Dec 1;52(6):783–9.

49. Loose M, Naber KG, Hu Y, Coates A, Wagenlehner FME. Urinary bactericidal activity of colistin and azidothymidine combinations against mcr-1-positive colistin-resistant Escherichia coli. Int J Antimicrob Agents. 2019 Jul;54(1):55–61.

50. Buckner MMC, Laura Ciusa M, Meek RW, Moorey AR, McCallum GE, Prentice EL, et al. HIV drugs inhibit transfer of plasmids carrying extended-spectrum β-lactamase and carbapenemase genes. MBio. 2020 Feb 25;11(1):e03355–19.

51. Peyclit L, Khedher M Ben, Zerrouki L, Diene SM, Baron SA, Rolain JM. Inactivation of thymidine kinase as a cause of resistance to zidovudine in clinical isolates of Escherichia coli: A phenotypic and genomic study. J Antimicrob Chemother. 2020;75(6):1410–4.

52. Doléans-Jordheim A, Bergeron E, Bereyziat F, Ben-Larbi S, Dumitrescu O, Mazoyer MA, et al. Zidovudine (AZT) has a bactericidal effect on enterobacteria and induces genetic modifications in resistant strains. Eur J Clin Microbiol Infect Dis. 2011;30(10):1249–56.

53. Maziarz EK, Perfect JR. Cryptococcosis. Infect Dis Clin North Am. 2016 Mar;30(1):179–206.

