## Supplemental data for "Zidovudine multi-combos with last-line fosfomycin, ceftazidime-avibactam, colistin and tigecycline against Multi-Drug Resistant *Klebsiella pneumoniae*"

Gómara-Lomero, M. et al.

**SUPPLEMENTARY FIGURES**

|  |  |  |  | **MIC (mg/L)** | | | | | | |
| --- | --- | --- | --- | --- | --- | --- | --- | --- | --- | --- |
| **Isolate** | **Resistance mechanism** | **Source** | **MDR/XDR** | **CST** | **FOF** | **TGC** | **ETP** | **MEM** | **CAZ-AVI** | **ZDV** |
| E-1 | CTX-M 14 | Rectal swab | XDR | 0.5 (S) | >64 (R) | 4 (R) | >32 (R) | 8 (I) | 1 (S) | 0.25-0.5 |
| E-2 | CTX-M 15 | Blood | MDR | 0.5 (S) | >64 (R) | 0.5 (S) | 64 (R) | 4-8 (I) | 1 (S) | 0.5-1 |
| E-3 | CTX-M 15 | Abscess | MDR | 1-2 (S) | >64 (R) | 4 (R) | 16 (R) | 2-4 (I) | 1 (S) | 2 |
| E-4 | CTX-M 15 | Blood | MDR | 0.5 (S) | >64 (R) | 4 (R) | 1 (R) | 0.03 (S) | 0.5 (S) | 1 |
| E-5 | SHV-1 + porin loss | Blood | MDR | 0.5 (S) | 8 (S) | 0.5-1 (S) | 0.25 (S) | 0.03 (S) | 0.06-0.12 (S) | 0.5-1 |
| A-6 | AmpC ACT-1 | SEIMC CCS07 | MDR | ≤0.5 (S) | >64 (R) | 1-2 (R) | 4-8 (R) | 0.5 (S) | 0.5 (S) | 8-16 |
| C-7 | OXA-48 | Blood | MDR | 1 (S) | >64 (R) | 2 (R) | 8-16 (R) | 4 (I) | 0.5 (S) | 2 |
| CS-8 | Colistin R | Urine | MDR | 16 (R) | >64 (R) | 1 (R) | 0.5 (S) | 0.5-1 (S) | 0.5 (S) | 0.5 |
| CSE-9 | VIM-1 + CTX-M 15 + colistin R | SEIMC CCS04 | XDR | 16 (R) | >64(R) | 1-2 (R) | 8-16 (R) | 16-32 (R) | >64 (R) | ≥64 |
| CE-10 | CTX-M 15 + OXA-48 | Blood | MDR | 1-2 (S) | >64 (R) | 1-2 (R) | 8 (R) | 4 (I) | 0.25 (S) | 64 |
| CEE-11 | KPC-3 + SHV-11 + TEM-1 | SEIMC CCS05 | XDR | 2 (S) | >64 (R) | 4 (R) | >64 (R) | >64 (R) | 4 (S) | 0.5-1 |
| CSEE-12 | OXA-1 + SHV-1 + colistin R | EARS QC | MDR | 4 (R) | 64 (R) | 1 (R) | 8-16 (R) | 1-2 (S) | 0.5 (S) | 1 |

**Table S1.** **Strain characterization of K. pneumoniae isolates and susceptibility profile to drugs evaluated in this study.** Clinical categorization according to EUCAST breakpoints (26) are displayed in brackets. MIC values were obtained by broth microdilution method in CAMHB. For CAZ-AVI and FOF MIC determination, medium was supplemented with 4 mg/L of avibactam and 25 mg/L of glucose-6-phosphate respectively.

MDR: non-susceptible to ≥1 agent in ≥3 antimicrobial categories; XDR: non-susceptible to ≥1 agent in all but ≤2 categories (categorization according to susceptibility results provided in Table S2); CAZ-AVI, ceftazidime-avibactam; CST, colistin; FOF, fosfomycin; ETP, ertapenem; MEM, meropenem; TGC, tigecycline; ZDV, zidovudine

**Figure S1. Time-kill curves of pairwise combinations currently used in the therapy of MDR enterobacteria against twelve *K. pneumoniae* clinical strains**

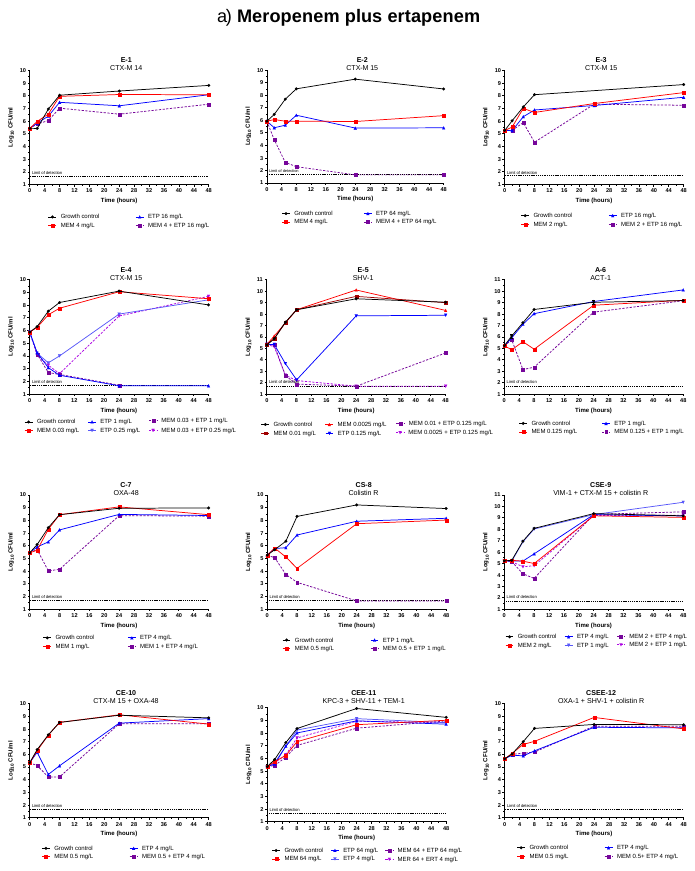

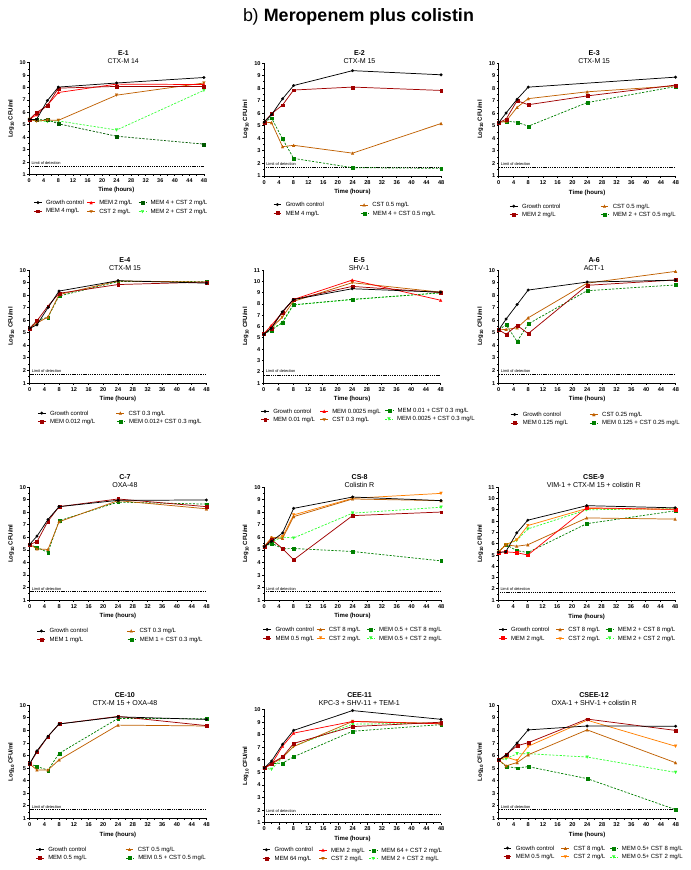

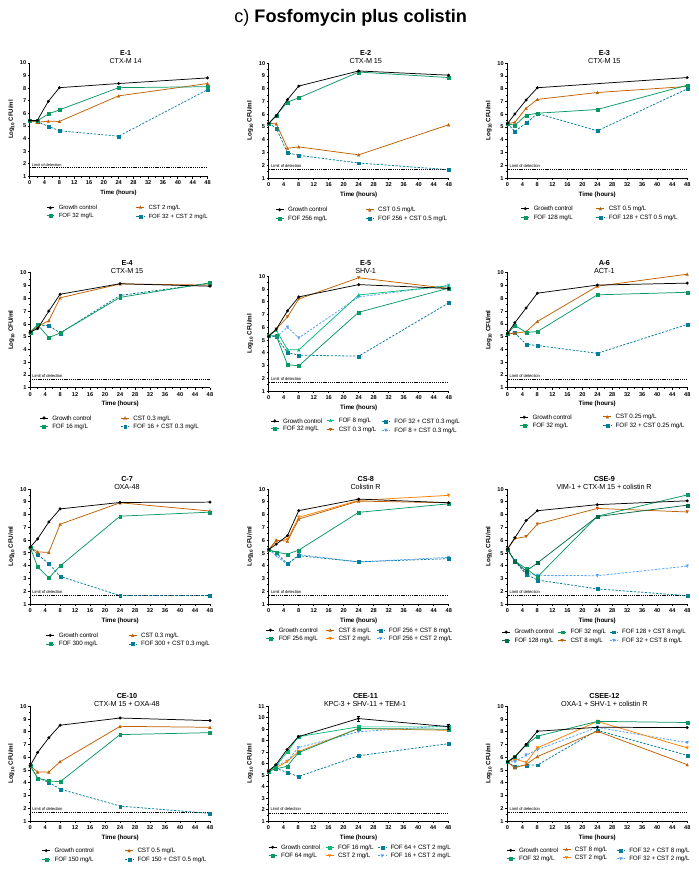

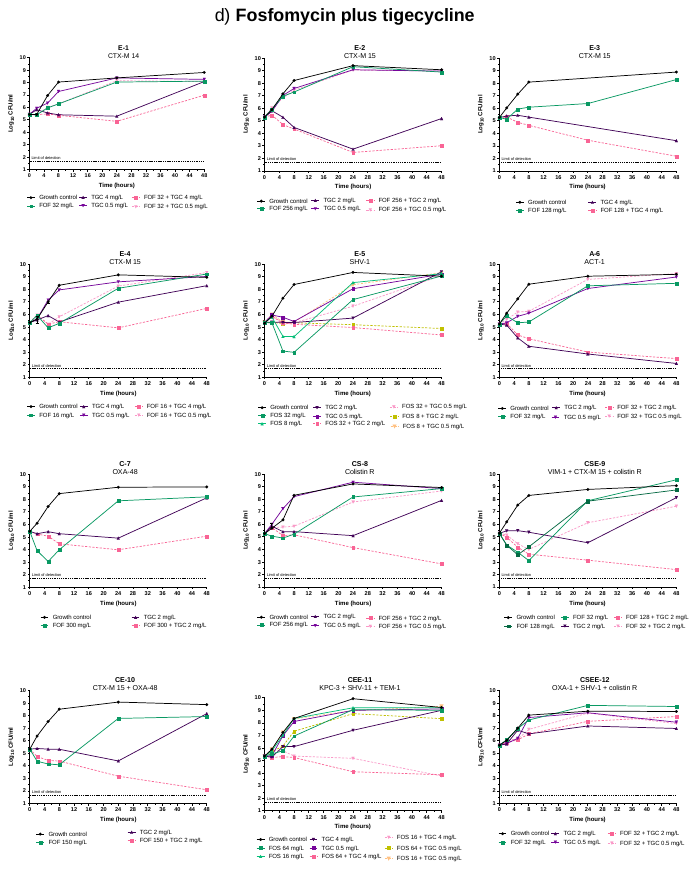

**Figure S2.** **Time-kill assays of zidovudine combined with last-line antibiotics against twelve *K. pneumoniae* clinical strains.**

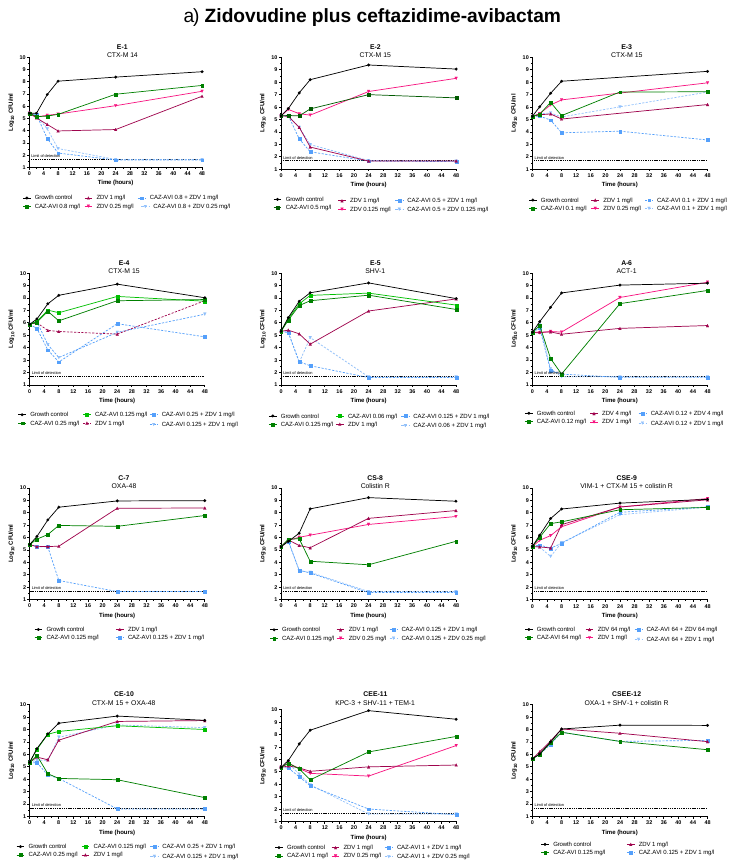

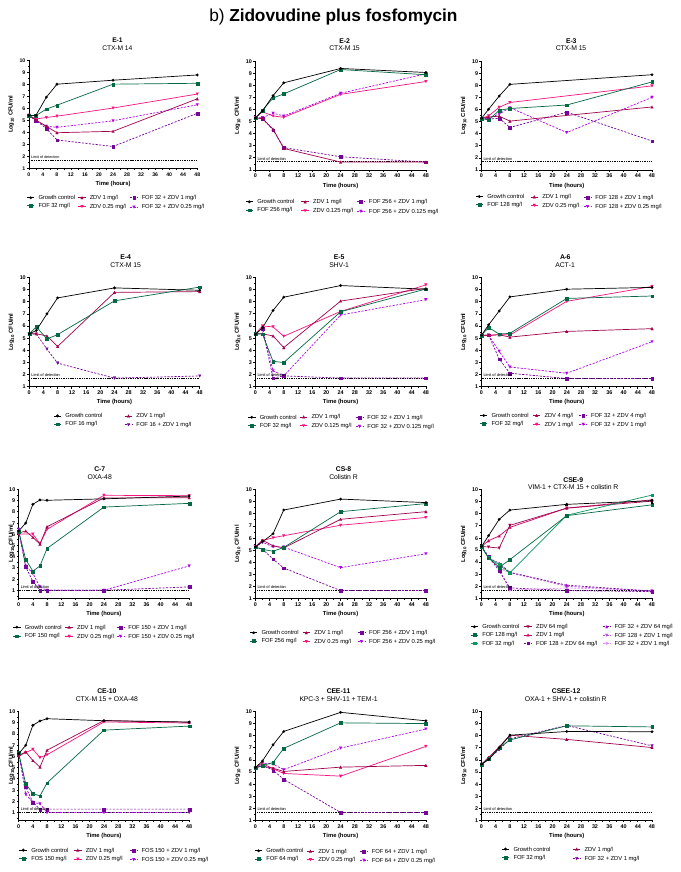

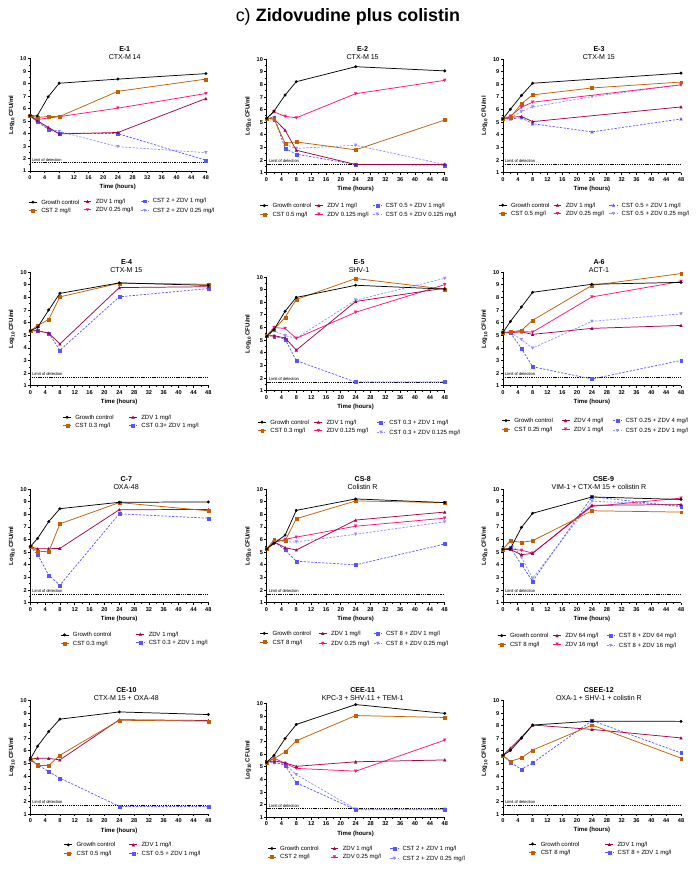

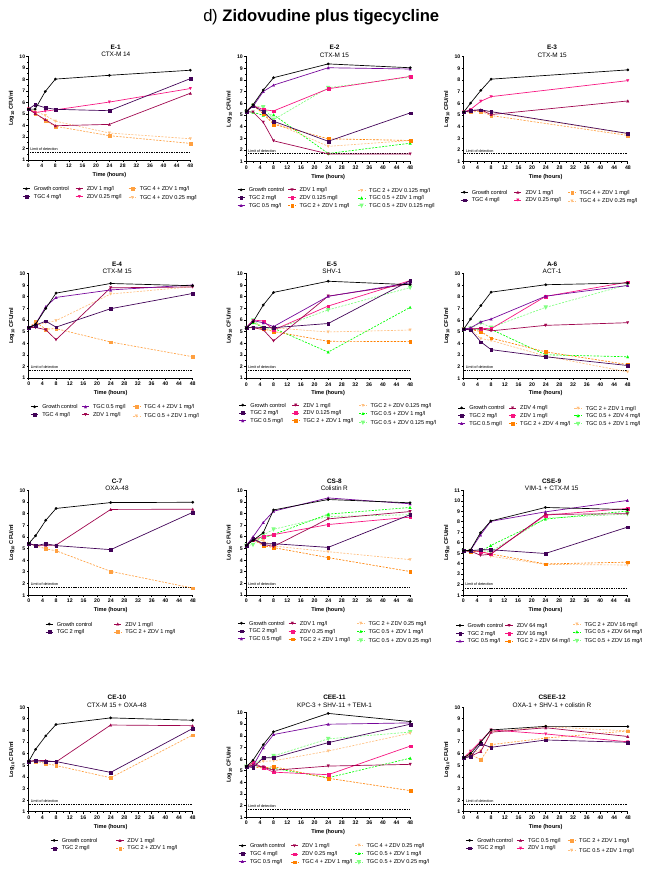
