## Supplemental Table S2 for "Zidovudine multi-combos with last-line fosfomycin, ceftazidime-avibactam, colistin and tigecycline against Multi-Drug Resistant *Klebsiella pneumoniae*"

**Table S2. Strain characterization and antimicrobial susceptibility provided by the Miguel Servet University Hospital (Zaragoza, Spain) for the twelve MDR/XDR K. pneumoniae isolates**

|  |  |  |  | **MIC (mg/L)^b^** | | | | | | | | | | | | | | | | | | | | |
| --- | --- | --- | --- | --- | --- | --- | --- | --- | --- | --- | --- | --- | --- | --- | --- | --- | --- | --- | --- | --- | --- | --- | --- | --- |
| **Isolate** | **Resistance mechanism** | **Specimen source** | **MDR/XDR classification^a^** | **AMK** | **GEN** | **TOB** | **AMP/AMX** | **AMC** | **TZP** | **FOX** | **CXM** | **CTX** | **CAZ** | **FEP** | **ATM** | **IPM** | **ETP** | **MEM** | **CIP** | **LVX** | **FOF** | **CST** | **TGC** | **SXT** |
| E-1 | CTX-M 14 | Rectal swab | XDR | ≤8 | >8 | ≤2 | >16 | >16/8 | >64 | >16 | >16 | >32 | >16 | >16 | >16 | 8 | >1 | 8 | >2 | 4 | ≤32 | ≤2 | >2 | >4/76 |
| E-2 | CTX-M 15 | Blood | MDR | ≤8 | ≤2 | >8 | >16 | >16/8 | >64 | >16 | >16 | >32 | >16 | >16 | >16 | ≤1 | >1 | 2 | >2 | >4 | >64 | ≤2 | ≤1 | ≤2/38 |
| E-3 | CTX-M 15 | Abscess | MDR | ≤8 | >8 | >8 | >16 | >16/8 | 64 | >16 | >16 | >32 | >16 | >16 | >16 | ≤1 | >1 | 2 | >2 | 2 | ≤32 | ≤2 | ≤1 | >4/76 |
| E-4 | CTX-M 15 | Blood | MDR | 16 | >4 | >4 | >16 | >32 | 16 | >16 | >8 | >32 | 32 | >8 | >4 | ≤1 | ≤0.12 | ≤0.12 | >1 | >1 | <=16 | ≤2 | >2 | >4/76 |
| E-5 | SHV-1 + porin loss | Blood | MDR | ≤8 | ≤2 | ≤2 | >16 | >16/8 | >64 | 16 | ≤4 | ≤1 | ≤1 | 4 | ≤1 | ≤1 | ≤0.5 | ≤1 | ≤0.5 | ≤1 | ≤32 | ≤2 | ≤1 | ≤2/38 |
| A-6 | AmpC ACT-1 | SEIMC CCS07 | MDR | >32 | >8 | >8 | >16 | >16/8 | 64 | >16 | >16 | 32 | >16 | ≤1 | >16 | ≤1 | >1 | ≤1 | 2 | ≤1 | ≤32 | ≤2 | ≤1 | >4/76 |
| C-7 | OXA-48 | Blood | MDR | ≤8 | ≤2 | ≤2 | >16 | >16/8 | >64 | ≤8 | 8 | ≤1 | ≤1 | ≤1 | ≤1 | 4 | >1 | 4 | ≤0.5 | ≤1 | 64 | ≤2 | ≤1 | ≤2/38 |
| CS-8 | Colistin R | Urine | MDR | ≤8 | ≤2 | ≤2 | >16 | >16/8 | >64 | >16 | 16 | ≤1 | ≤1 | 4 | ≤1 | ≤1 | ≤0.5 | ≤1 | ≤0.5 | ≤1 | >64 | >4 | 2 | >4/76 |
| CE-9 | VIM-1 + CTX-M 15 + colistin R | SEIMC CCS04 | XDR | 16 | >8 | >8 | >16 | >16/8 | >64 | >16 | >16 | >32 | >16 | >16 | >16 | 2 | >1 | 8 | >2 | >4 | ≤32 | >4 | 2 | >4/76 |
| CE-10 | CTX-M 15 + OXA-48 | Blood | MDR | ≤8 | >4 | >4 | >8 | >32 | >16 | ≤8 | >8 | >32 | 32 | >8 | >4 | 8 | >1 | 1 | >1 | >1 | 32 | ≤2 | ≤1 | >4/76 |
| CEE-11 | KPC-3 + SHV-11 + TEM-1 | SEIMC CCS05 | XDR | 32 | 4 | >8 | >16 | >16/8 | >64 | >16 | >16 | >32 | >16 | >16 | >16 | >8 | >1 | >8 | >2 | >4 | 64 | >4 | >2 | >4/76 |
| CSEE-12 | OXA-1 + SHV-1 + colistin R | EARS QC | MDR | >32 | >8 | >8 | >16 | >16/8 | >64 | ≤8 | >16 | >32 | ≤1 | >16 | ≤1 | ≤1 | >1 | ≤1 | >2 | >4 | ≤32 | >4 | 2 | >4/76 |

^a^MDR: non-susceptible to ≥1 agent in ≥3 antimicrobial categories; XDR: non-susceptible to ≥1 agent in all but ≤2 categories

^b^Automated MIC method (Microscan Walkaway®, Beckman Coulter, Spain)

| AMK | Amikacin |
| --- | --- |
| GEN | Gentamicin |
| TOB | Tobramicin |
| AMP/AMX | Ampicillin/amoxicillin |
| AMC | Amoxicillin-clavulanate |
| TZP | Piperacillin-tazobactam |
| FOX | Cefoxitin |
| CXM | Cefuroxime |
| CTX | Cefotaxime |
| CAZ | Ceftazidime |
| FEP | Cefepime |
| ATM | Aztreonam |
| IPM | Imipenem |
| ETP | Ertapenem |
| MEM | Meropenem |
| CIP | Ciprofloxacin |
| LVX | Levofloxacin |
| FOF | Fosfomycin |
| CST | Colistin |
| TGC | Tigecycline |
| SXT | Trimethoprim/sulfamethoxazole |
